# Integrating genomics into clinical pediatric oncology using the molecular tumor board at the Memorial Sloan Kettering Cancer Center

**DOI:** 10.1101/040584

**Authors:** Michael V. Ortiz, Rachel Kobos, Michael Walsh, Emily K. Slotkin, Stephen Roberts, Michael F. Berger, Meera Hameed, David Solit, Marc Ladanyi, Neerav Shukla, Alex Kentsis

**Affiliations:** Department of Pediatrics, Sloan Kettering Institute, Memorial Sloan Kettering Cancer Center, New York, NY; Department of Medicine, Sloan Kettering Institute, Memorial Sloan Kettering Cancer Center, New York, NY; Department of Pathology, Sloan Kettering Institute, Memorial Sloan Kettering Cancer Center, New York, NY; Human Oncology and Pathogenesis Program, Sloan Kettering Institute, Memorial Sloan Kettering Cancer Center, New York, NY; Molecular Pharmacology Program, Sloan Kettering Institute, Memorial Sloan Kettering Cancer Center, New York, NY

**Keywords:** Pediatric oncology, Molecular biology, New agents, Cancer pharmacology

## Abstract

**Background:** Pediatric oncologists have begun to leverage tumor genetic profiling to match patients with targeted therapies. At the Memorial Sloan Kettering Cancer Center (MSKCC), we developed the Pediatric Molecular Tumor Board (PMTB) to track, integrate, and interpret clinical genomic profiling and potential targeted therapeutic recommendations.

**Procedure:** This retrospective case series includes all patients reviewed by the MSKCC PMTB from July 2014 to June 2015. Cases were submitted by treating oncologists and potential treatment recommendations were based upon the modified guidelines of the Oxford Centre for Evidence Based Medicine.

**Results:** There were 41 presentations of 39 individual patients during the study period. Gliomas, acute myeloid leukemia, and neuroblastoma were the most commonly reviewed cases. Thirty nine (87%) of the 45 molecular sequencing profiles utilized hybrid-capture targeted genome sequencing. In 30 (73%) of the 41 presentations, the PMTB provided therapeutic recommendations, of which 19 (46%) were implemented. Twenty-one (70%) of the recommendations involved targeted therapies. Three (14%) targeted therapy recommendations had published evidence to support the proposed recommendations (evidence levels 1-2), 8 (36%) recommendations had preclinical evidence (level 3), and 11 (50%) recommendations were based upon hypothetical biological rationales (level 4).

**Conclusions:** The MSKCC PMTB enabled a clinically relevant interpretation of genomic profiling. Effective use of clinical genomics is anticipated to require new and improved tools to ascribe pathogenic significance and therapeutic actionability. Development of specific rule-driven clinical protocols will be needed for the incorporation and evaluation of genomic and molecular profiling in interventional prospective clinical trials.

MSKCC: Memorial Sloan Kettering Cancer Center
PMTB: Pediatric Molecular Tumor Board
ALL: Acute lymphoblastic leukemia
DSRCT: Desmoplastic small round cell tumor
AML: Acute myeloid leukemia

## Introduction

The recognition of cancer as a genetic disease prompted the incorporation of genetic profiling into clinical care to improve the diagnostic accuracy and stratification of conventional therapies. For example, diagnosis and therapy stratification of childhood acute lymphoblastic leukemia (ALL) based on ploidy and chromosomal rearrangements have substantially improved the long-term survival for this group of patients. [1] Discovery of genetic alterations that confer susceptibility to specifically targeted therapies, such as tyrosine kinase inhibitors and retinoic acid for *BCR-ABL1* and *PML-RARα* rearranged leukemias, respectively, have enabled prospective identification of patients who would benefit from rationally targeted therapies, leading to transformative improvements in their outcomes. [1, 2] In addition, genetic alterations have been used as specific diagnostic markers, such as *EWS-FLI1* and *EWS-WT1* in Ewing sarcoma and desmoplastic small round cell tumor (DSRCT), respectively. [3]

Given the increasing feasibility of genome profiling and number of clinically available targeted therapies [4, 5], academic medical centers have begun to deploy multiplexed genomic profiling assays to match patients with investigational or approved targeted agents. For example, Tsimberidou and colleagues used genomic profiling in assignment of patients to phase 1 clinical trials based on the identification of tumor mutations that may confer susceptibility to relevant investigational drugs. [6] Recently, Mody *et al* and Beltran *et al* used exome sequencing of tumors of patients with relapsed or refractory disease to identify potential therapeutically actionable lesions, leading to alterations in therapy in subsets of patients. [7, 8] Notably, Rubio-Perez and colleagues found that although only 5.9% of tumors in their cohort had mutations that were potentially susceptible to approved drugs, up to 73% of tumors may be susceptible to drugs that are currently under investigation or are already approved for other indications. [9]

As a result of these and other studies, two features of current cancer genome profiles have emerged to inform the integration of molecular profiling into clinical oncology: i) many known cancer-causing mutations in individual tumors tend to occur at relatively low frequencies in large unselected patient cohorts, and ii) only a minority of observed gene alterations implicated in cancer pathogenesis can be ascribed a pathogenic function and approved therapeutic agent at the present time. In addition, it is not yet known whether incorporation of genomic profiling into routine clinical care, particularly for patients with relatively rare cancer types such as children, will lead to improvements in clinical outcomes.

To enable the long-term investigation of these questions, we have developed a Pediatric Molecular Tumor Board (PMTB) to track, integrate, and offer potential therapeutic recommendations based on clinical genomic tumor profiling at the Memorial Sloan Kettering Cancer Center (MSKCC). Here, we report our experience during the first year of this program, and discuss implications for the effective integration of molecular profiling into clinical pediatric oncology.

## Methods

### Study Design

This is a retrospective case series of all patients reviewed at the MSKCC PMTB from July of 2014 to June of 2015. MSKCC is a tertiary academic medical center, caring for both local and referred patients.

### Molecular Profiling

All histopathologic and molecular data obtained as part of routine clinical care were included. All patients were consented and enrolled on institutionally approved tissue specimen acquisition and molecular profiling protocols. We obtained multiplexed genomic assays from MSK-IMPACT, a hybrid capture-based DNA sequencing assay of 341 or 410 genes, depending on the utilized assay version [10, 11], FoundationONE Heme, a hybrid capture-based DNA and RNA sequencing assay targeting 405 genes involved in hematologic malignancies [12], whole-exome sequencing [8], and a 30-gene panel of recurrently mutated genes in myeloid malignancies [13]. Analysis of constitutional or germ-line mutations and pathogenic alleles was explicitly included in the informed consent process, and depended on case-by-case review by a dedicated clinical pediatric geneticist for the interpretation of potential pathogenicity and return of information to patients.

### Pediatric Molecular Tumor Board

Cases for monthly PMTB review were submitted by the primary oncology physicians at MSKCC based on their own assessment of the need for PMTB review. Referring physicians provided summaries of relevant clinical information to the PMTB organizers, who reviewed the pertinent clinical, pathological, and molecular profiling data before each PMTB meeting. The PMTB was comprised of pediatric oncologists, pathologists, geneticists, bioinformatics specialists, and cancer biologists with relevant expertise based on specific cases presented.

Annotation of known pathogenic mutations was provided by the respective clinical genomics assays, as part of their standardized mutation calling. [8, 11, 12] For mutations without reported annotation and for variants of unknown significance, interpretation included individual review of the published literature and cancer genome databases, including canSAR [14], cBioPortal for Cancer Genomics [15, 16], and Tumor Portal [17]. Novel missense gene mutations were modeled using structure-homology modeling with SWISS-MODEL. [18] Functional significance of observed mutations and variants was assessed based on tumor purity from histopathologic and DNA sequencing data, detected allele frequency, and functional assessments based on published literature and biological predictions, as synthesized by the PMTB review.

Treatment recommendations were made using modified guidelines of the Oxford Centre for Evidence Based Medicine: drug approved for specific indication with known pathogenic mutation (level 1), clinical evidence supporting the ‘off-label’ use of an approved drug (level 2), preclinical evidence demonstrating benefit (level 3), and mechanism-based rationale without direct preclinical evidence of efficacy (level 4). [19] Case assessments, profile interpretation, and clinical recommendations were recorded and disseminated using a dedicated, Health Insurance Portability and Accountability Act compliant, access-controlled online database. Level 3 and 4 recommendations were made only for patients for whom standard therapy failed and who were deemed not eligible for active clinical trials.

## Results

During the one-year course of this study, 41 cases of 39 individual patients were reviewed in the course of 11 monthly PMTB sessions. **Table I** describes the specific features of the PMTB patient cohort as compared to all pediatric patients treated at MSKCC during the same period of time. There were 2-7 (median = 3) presentations per PMTB meeting. The median age and gender distribution of PMTB-reviewed patients were 13 years and 67% male, respectively, as compared to 11 years and 57% male for all patients treated in the Department of Pediatrics during the same period of time. Four patients in the study cohort were over 21 years of age based on the referral to pediatric oncologists due to the diagnosis of primarily pediatric cancers, e.g. neuroblastoma. The most common malignancies in the reviewed cohort included high-grade glioma (20%), acute myeloid leukemia (AML) (15%), and neuroblastoma (13%), as shown in **Figure 1**. These diagnoses represented 2%, 6%, and 18% of the overall pediatrics cohort, respectively. In contrast, the three most common malignancies treated for all pediatric oncology patients during the same time period at MSKCC were neuroblastoma (18%), sarcomas excluding rhabdomyosarcoma (18%), and acute lymphoblastic leukemia (ALL) (11%). Three quarters of the patients presented at PMTB had either relapsed or refractory disease. Some cases involved patients in remission based on the high-risk or unusual nature of their disease, as assessed by their primary oncologists.

**Table I:**
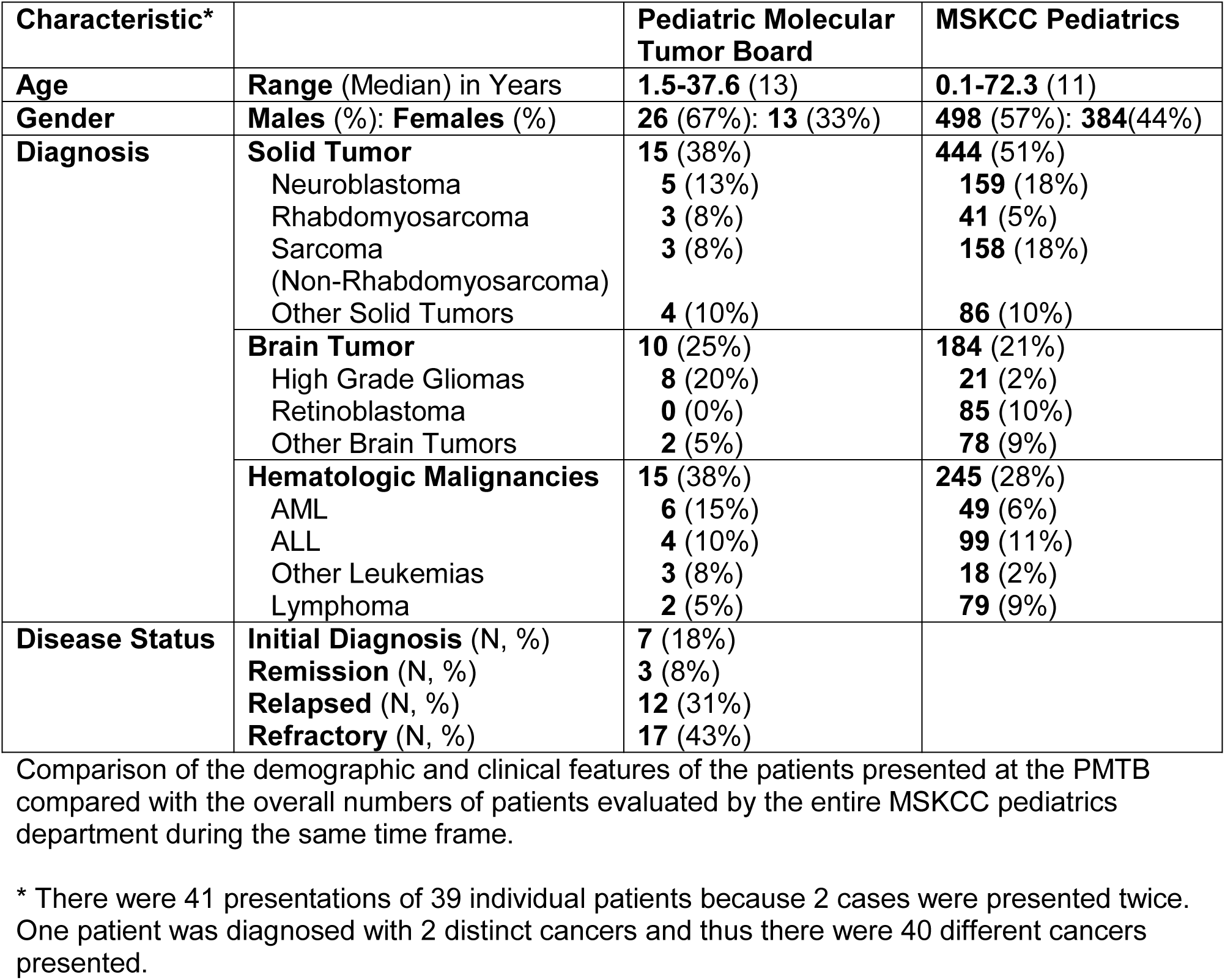
Characteristics of patients presented at the Pediatric Molecular Tumor Board.

**Figure 1.**
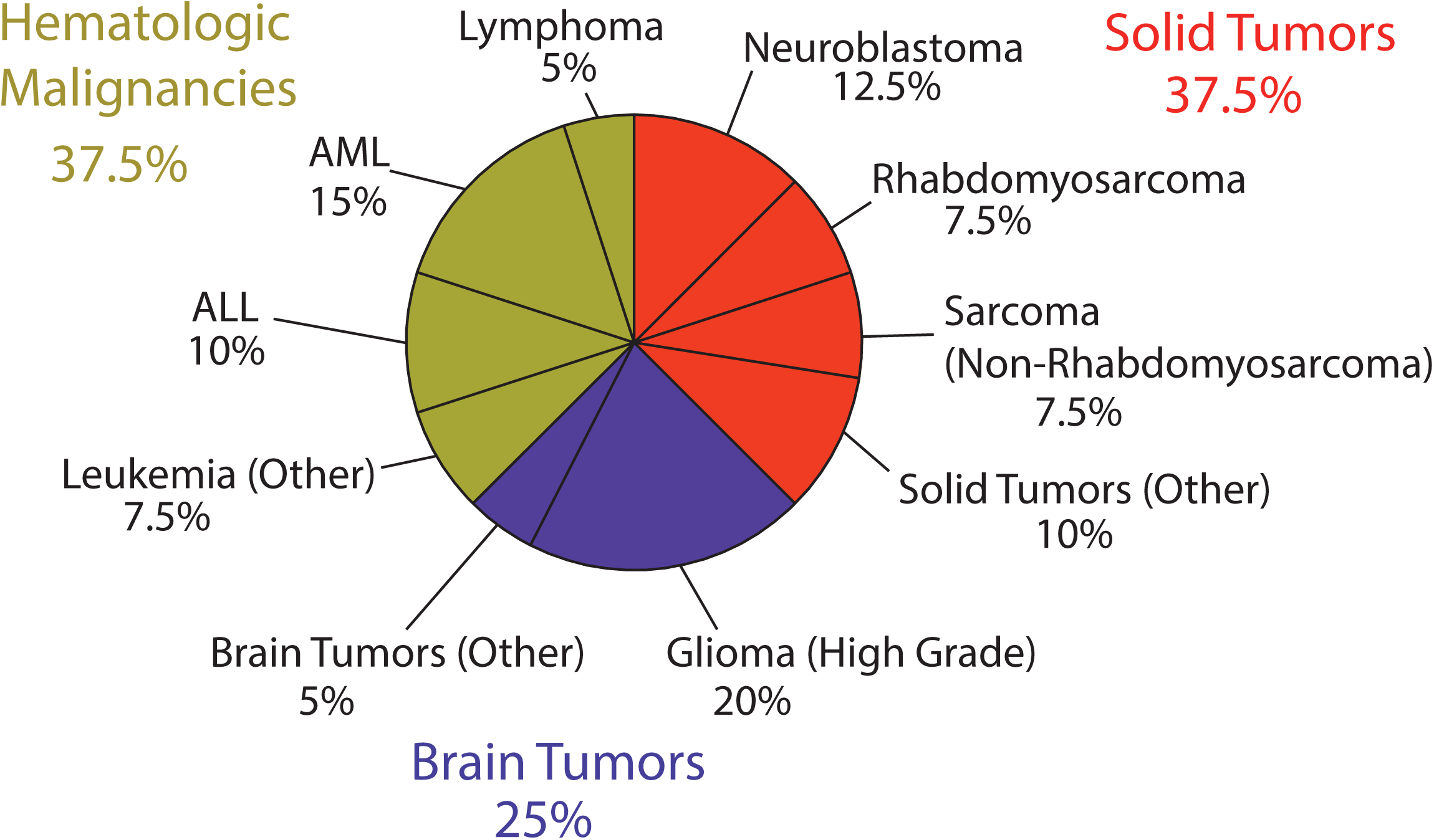
Distribution of PMTB oncologic diagnoses. PMTB reviewed molecular profiling results for hematologic (green), solid (red) and brain (blue) tumors. High-grade gliomas, acute myeloid leukemia, and neuroblastoma were the most common diagnoses presented to the PMTB.

For each PMTB, the primary referring oncologists had ordered tumor molecular profiling as per their individual assessments, and submitted them for PMTB review. Reviewed molecular profiles predominantly involved multiplexed gene panel sequencing (**Figure 2**). There were 45 total molecular profiles reviewed, of which 20 used MSK-IMPACT and 18 utilized FoundationONE Heme gene sequencing panels. Two cases underwent exome sequencing. Six cases discussed during the PMTB included cytogenetic or FISH assays.

**Figure 2.**
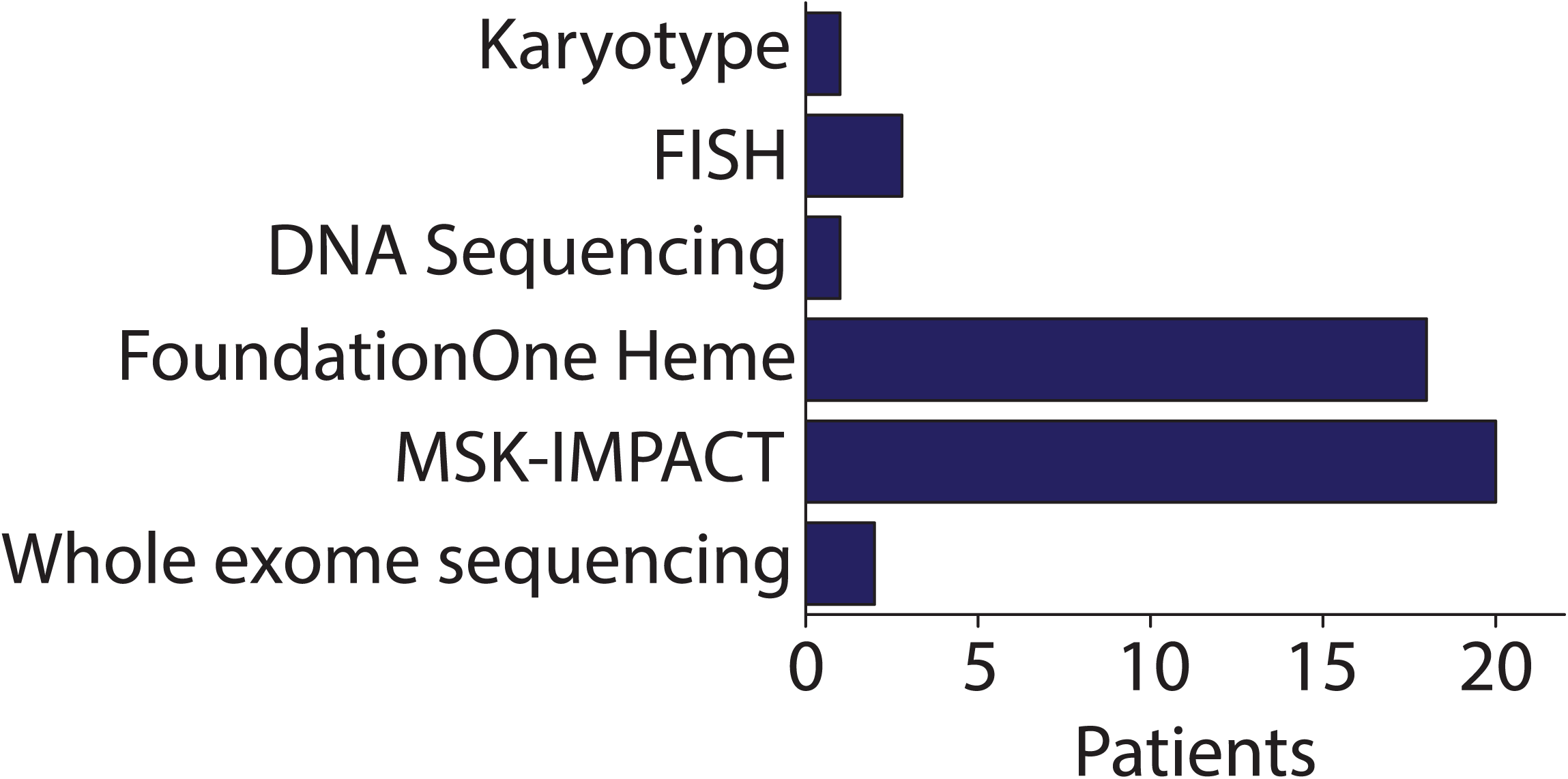
Distribution of PMTB cancer molecular profiling platforms. PMTB review involved cytogenetic and DNA sequencing results, including targeted and whole-exome sequencing. Majority of reviewed profiles involved targeted genomic profiling, such as FoundationOne Heme and MSK-IMPACT.

In total, there was at least one molecular aberration identified for every case with 3.9 gene alterations per tumor on average (median = 3), as shown in **Figure 3**. We reviewed 2 cases in which molecular profiling revealed known pathogenic mutations with approved targeted agents. For a patient with medullary thyroid carcinoma, molecular profiling identified an activating *RET* mutation (exon 11 p.D631_L633delinsE; c.1893_1899delinsA), prompting a recommendation for therapy with the tyrosine kinase inhibitor cabozantinib, based on the phase 3 study that demonstrated a statistically significant improvement in survival from 4 to 11 months when compared to placebo. [20, 21] For a patient with a diagnosis of Ewing sarcoma based on the apparent focal *EWSR1* rearrangement determined using FISH, genomic profiling instead demonstrated a complex genomic profile involving deletion of *RB1* and the absence of known pathogenic *EWSR1* rearrangements. This was found to be consistent with the diagnosis of osteosarcoma, which was confirmed histopathologically upon surgical resection post-neoadjuvant therapy. As a result, PMTB recommended to alter treatment to osteosarcoma directed platinum-based therapy. [22]

**Figure 3.**
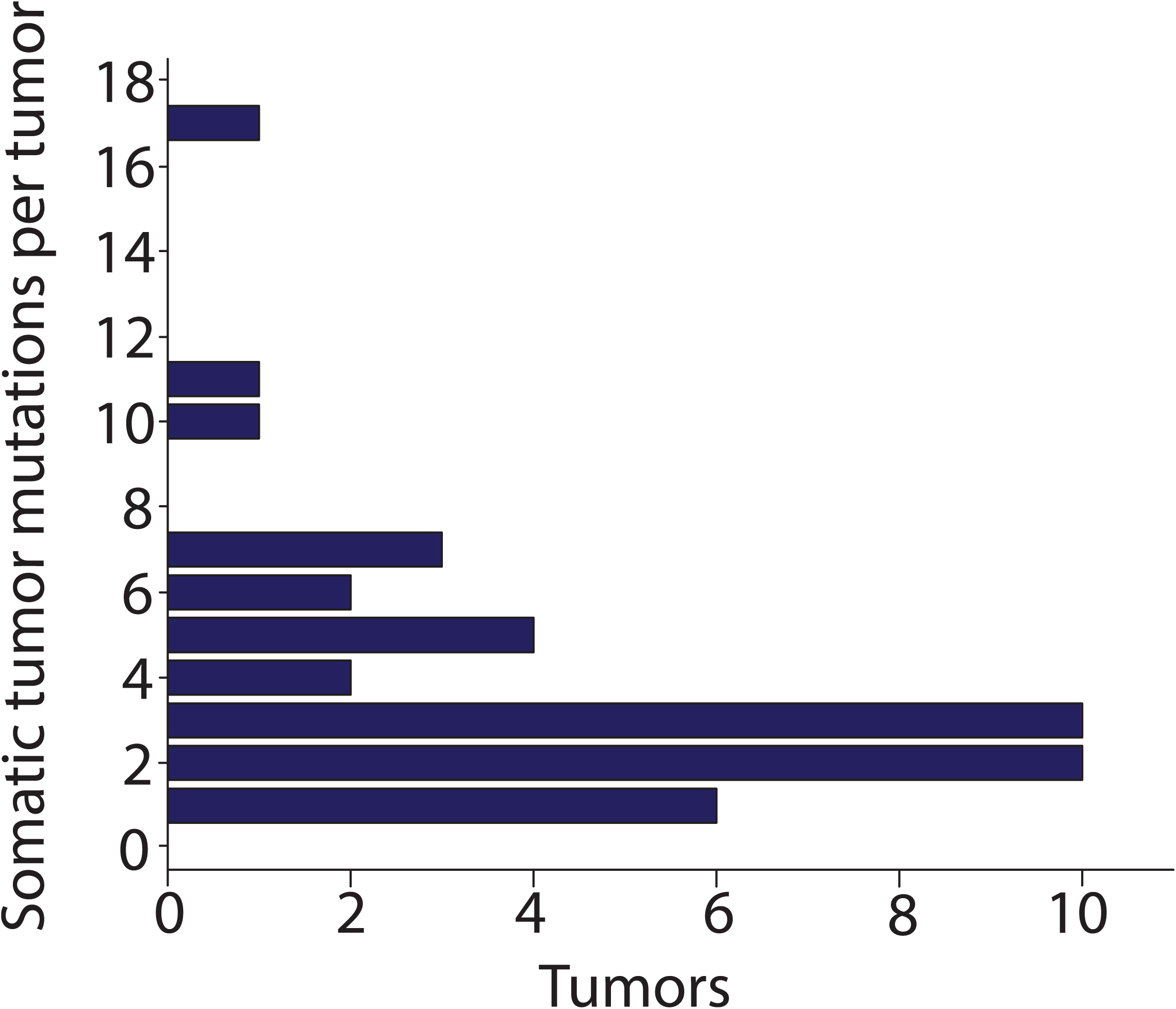
Distribution of the identified somatic tumor mutations. The median number of somatic mutations was found to be 3 per tumor. Over 90% of tumors had 1-7 somatic mutations.

For cases without standard-of-care therapies or those lacking approved therapies, modified Oxford Centre for Evidence Based Medicine guidelines were utilized to interpret observed genetic alterations and make potential clinical recommendations (see Methods for description of the modifications). [19] In total, PMTB made 30 recommendations, 24 of which were for alterations in therapy (**Figure 4**). For example, targeted RNA capture revealed *ZMIZ1-ABL1* fusion in a case of acute lymphoblastic leukemia, leading to the diagnosis of Ph-like ALL and recommendation for ‘off-label’ treatment with the tyrosine kinase inhibitor dasatinib based on the prediction that the proline rich-domain of ZMIZ1 mediates protein-protein interactions causing constitutive dimerization and ABL1 kinase activation (level 2). [23, 24]

**Figure 4.**
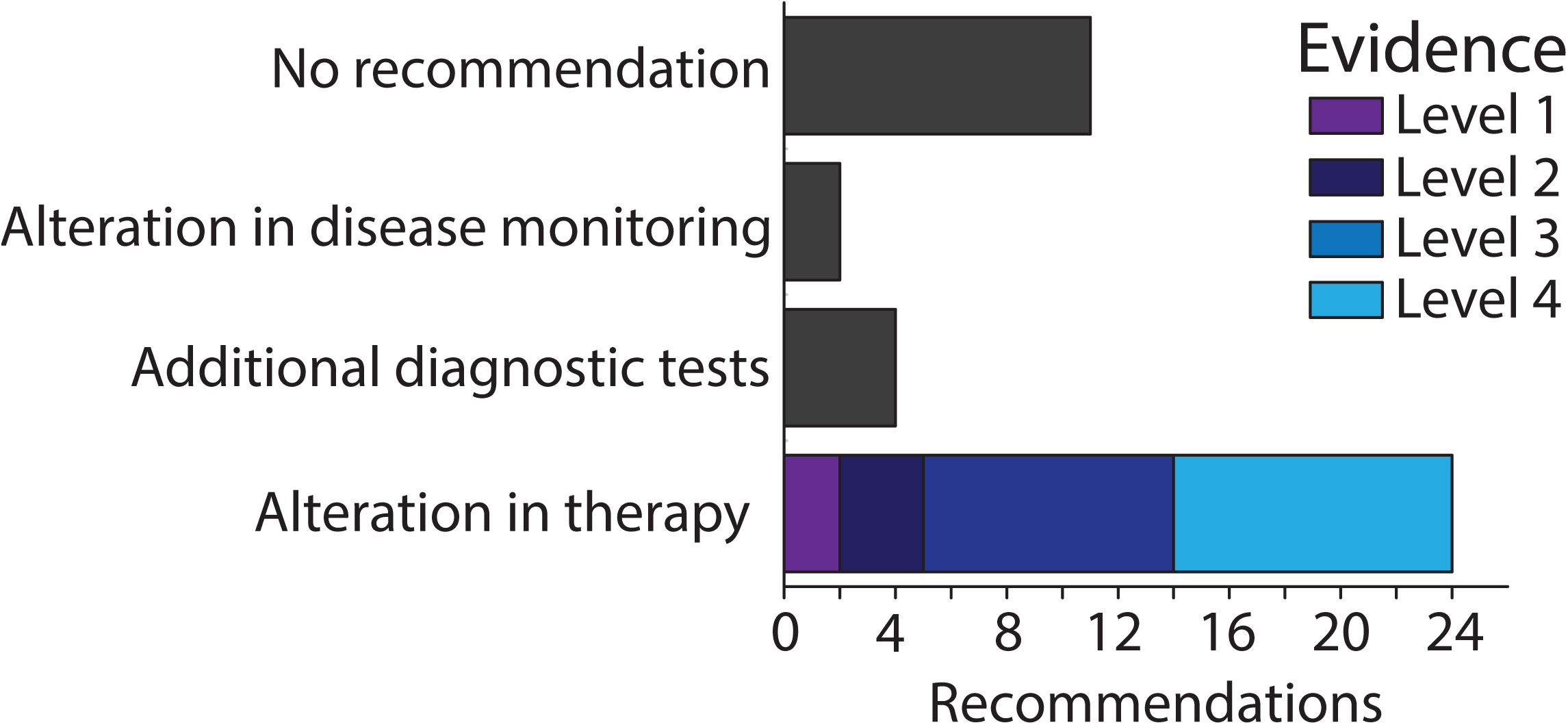
Recommendations of the PMTB. PMTB successfully provided recommendations based on molecular profiling results in the majority of the reviewed cases. Only a minority of cases for which alterations in therapy were made had clinical evidence to support therapeutic recommendations (levels 1-2).

We found that the majority of cases reviewed by the PMTB lacked published evidence of clinical efficacy of potential targeted therapies. Consequently, the majority of recommendations for alterations in treatment were based upon either preclinical evidence (level 3) or clinically hypothetical rationales based on biological evidence and inferred molecular mechanisms (level 4). To make treatment recommendations for cases lacking clinical evidence of therapeutic efficacy, we first ascertained whether a particular genetic alteration was likely to be pathogenic. For example, we identified a somatic non-sense tumor mutation of *PTCH1* (P25fs*54) in a case of neuroblastoma, which was previously reported as a pathogenic allele in medulloblastoma. As a result, we recommended potential therapy with vismodegib, based on the evidence that Smoothened (SMO) receptor inhibition is effective for Hedgehog signaling pathway-driven medulloblastomas and basal cell carcinomas. [25, 26] In contrast, vismodegib therapy was not recommended for a patient with relapsed mesenchymal chondrosarcoma and somatic heterozygous missense *PTCH1* mutation (exon 16 p.E864G; c.2591A>G), because of its uncertain likelihood of PTCH1 inactivation. Instead, additional testing for evidence of the functional activation of the Hedgehog signaling pathway, and potential susceptibility and consideration of SMO receptor inhibition, was recommended using GLI1 and GLI2 immunohistochemical testing. [27]

Once sufficient evidence of likely pathogenicity was obtained, PMTB sought to identify available therapies with potential efficacy. For example, a patient with hepatoblastoma with a somatic activating *CTNNB1* mutation (exon3 p.W25_H36del) was recommended therapy with the tyrosine kinase dasatinib based on the preclinical evidence that dasatinib inhibits activation of the YAP1 transcriptional complex that is required for survival of cells driven by activating *CTNNB1* mutations. [28]

We observed several cases for which multiple treatment options were available for a single potentially pathogenic mutation, necessitating the prioritization of the best approach. For example, a patient with refractory neuroblastoma and activating somatic *KRAS* A146T mutation was considered for potential therapy with either RAF or MEK kinase inhibitors. Given that RAF inhibitors are effective in blocking the signaling activity of *RAS* mutations but can be subject to resistance due to adaptive signaling, treatment with the MEK inhibitor trametenib that does not cause feedback resistance was recommended. [29, 30] This recommendation was further supported by the preclinical studies of neuroblastoma cell lines with constitutively active RAS signaling. [31-33] A similar rationale was developed for trametenib therapy for a patient with an anaplastic pleomorphic xanthoastrocytoma and potentially RAF-activating *BRAF*-*CCDC6* gene fusion that was recently reported to be associated with xanthoastrocytomas. [20, 34]

For some cases, we observed multiple potentially pathogenic mutations with available therapies, requiring prioritization of pathogenic alleles based on biological and therapeutic considerations. For example, a patient with AML was found to have both somatic rearrangement of *MLL* and *PTPN11* A72T and T507K mutations. Somatic *PTPN11* mutations are thought to be secondary mutational events in pediatric AML, and appear to have no prognostic significance. [35, 36] Given the essential oncogenic activity of *MLL* fusion genes and current preclinical evidence of DOT1L methyltransferase inhibition in the treatment of MLL-rearranged leukemias [37], recommendation for potential enrollment on the NCT02141828 clinical trial of the DOT1L inhibitor EPZ-5676 was made. Similarly, we recommended therapy with the MEK inhibitor trametenib for a patient with glioblastoma multiforme and somatic mutations of *NF1* (exon9 p.R304X; c.910C>T) and *FGFR1* (exon 13 p.N577K; c.1731C>G), which are both expected to cause activation of RAS-RAF-MEK signaling. [38]

Although most of the sequencing analyses focused on the identification of somatic pathogenic mutations with therapeutic implications, in certain cases PMTB recommendations were also based on the findings of constitutional or germ-line mutations. For example, a patient with AML and history of osteosarcoma was found to have a germ-line inactivating homozygous *PMS2* mutation, 1687C>T (R563X). [39] Given this genetic mutation consistent with a constitutional mismatch repair defect, the PMTB recommended treatment with the PD-1 receptor checkpoint inhibitor pembrolizumab based on the recent evidence that mismatch repair-deficient tumors may be susceptible to immune checkpoint blockade due to the increased presentation of neo-antigenic epitopes. [40] The interpretation and return of results of constitutional and germ-line mutational analysis were conducted in consultation with a dedicated pediatric clinical cancer geneticist, including referrals for family genetic counseling.

In summary, PMTB review led to 21 recommendations of targeted therapeutic agents, as shown in **Figure 5**. Three (14%) targeted therapy recommendations had clinical evidence to support the proposed recommendations (evidence levels 1-2), 8 (36%) recommendations had preclinical evidence (level 3), and 11 cases (50%) were based upon mechanism-based reasoning (level 4). Decision to offer PMTB recommended therapies were at the discretion of the primary treating oncologists. In retrospective review, we found that 15 of the 24 PMTB-recommended therapies were prescribed and administered in concordance with the PMTB review. We found diverse causes for the nine patients who did not receive PMTB-recommended therapies. Two patients died of progressive disease before being able to receive recommended therapies. In three cases, the primary oncologists elected to delay the implementation of treatment recommendations until possible disease progression. In one case, a patient with adenoid cystic carcinoma and somatic inactivating *ARID1A* mutation (exon 1 p. S11fs; c.31_56del) was recommended therapy with the EZH2 methyltransferase inhibitor EPZ-6438, based on the potential therapeutic efficacy of EZH2 inhibition in ARID1A-deficient tumors. [41] Because of the eligibility restrictions of the NCT01897571 EPZ-6438 clinical trial, the patient was instead treated with regorafenib as part of the NCT02098538 clinical trial. [42, 43] Finally, two patients were lost to follow up and could not be assessed. The majority of cases reviewed in PMTB involved patients with relapsed or refractory disease after the failure of standard-of-care therapies. As a result, considerations of active clinical trials tended to involve potential phase 1 and 2 studies.

**Figure 5.**
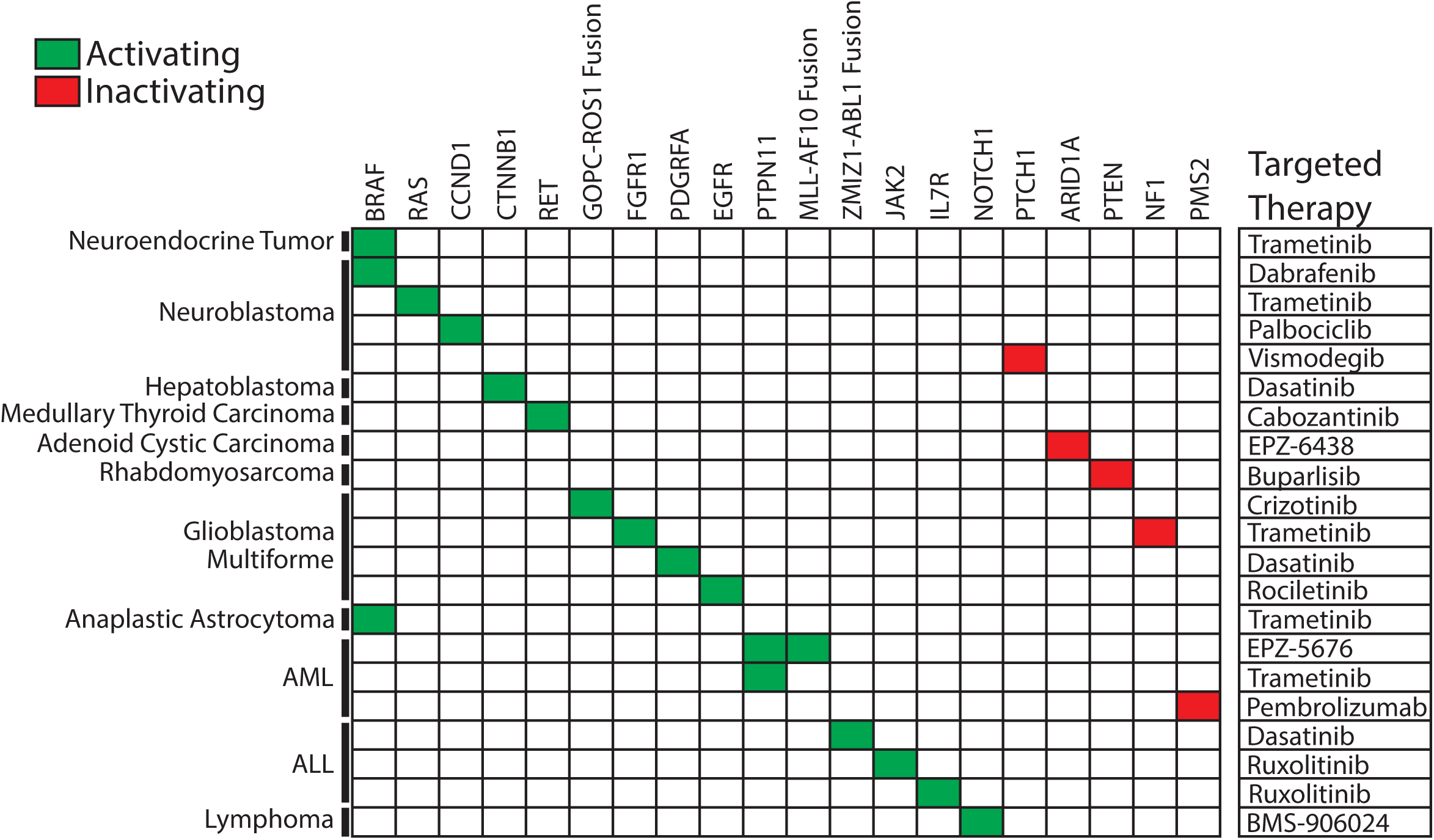
Molecular targets and therapies recommended by the PMTB. PMTB made recommendations of targeted therapies (right column) for both activating (green) and inactivating (red) gene mutations.

## Discussion

Based on the increasing use of clinical genomics and molecular profiling, we implemented the Pediatric Molecular Tumor Board at the Memorial Sloan Kettering Cancer Center. Its voluntary participation by treating oncologists led to review of a significant number of patients that comprised a minority of all pediatric patients treated at MSKCC during the same period of time (Table I). PMTB was able to provide therapeutic interpretations of molecular profiles for the majority of patients (Figures 3 & 4), and targeted therapies were recommended for 21 of the cases (Figure 5). We found that only a handful of current molecular profiles involved aberrations with level 1 evidence supporting specific therapeutic recommendations (Figure 2). Consequently, PMTB’s ability to offer clinically useful interpretations of current molecular profiles required: i) synthesis of published evidence of the prevalence of observed alleles and their documented pathogenicity, ii) inference of potential pathogenicity based on molecular and signaling pathway modeling, and iii) inference of potential therapeutic susceptibility based on the apparent allelic frequencies of observed mutations and known drug mechanisms of action.

For example, we made recommendations for targeted therapy of somatic cancer alleles in a particular tumor type when functionally similar alleles have been reported to be pathogenic in other tumor types, e.g. SMO inhibitor vismodegib of inactivating nonsense *PTCH1* allele in neuroblastoma. In contrast, mis-sense *PTCH1* mutation with low likelihood of functional dysregulation based on molecular modeling instead led to recommendation for additional evidence of functional activity (GLI1 and GLI2 immunohistochemistry). Likewise, we made treatment recommendations when observed alleles were predicted to be pathogenic based on molecular modeling, e.g. novel *ABL1* gene fusion and JAK1 F779L mutation that were predicted to cause constitutive kinase activation with potential susceptibility to tyrosine kinase inhibition (TKI) in Ph-like ALL and neuroblastoma, respectively. We avoided making therapeutic recommendations based on mutations with low allelic frequencies that suggested their sub-clonal origins, instead prioritizing apparently clonal mutations with high allelic frequencies as essential and pathogenic. Finally, we explicitly took into account potential for therapeutic efficacy and resistance, prioritizing molecular targets and drugs with no immediate or known resistance mechanisms, e.g. the use of MEK as opposed to RAF inhibitors in *BRAF*-mutant cancers. Similarly, we recommended against therapy with erlotinib or gefitinib for a patient with EGFR H773 insertion (c.2317 2319dup) mutant glioblastoma because of the inherent resistance of this kinase mutation to ATP-competitive kinase inhibitors [44], instead recommending potential therapy with third-generation EGFR-directed TKIs such as rociletinib once they become available for pediatrics and demonstrate activity against this mutation type. [45, 46]

Although we found that the PMTB provided clinical decision support and therapeutic recommendations based on genomic and molecular profiles, we encountered several significant challenges. First, we observed that the specific molecular profiling technologies, chosen by the treating oncologists based on their own considerations, influenced our ability to make treatment recommendations. Principally, this was driven by the choice between highly sensitive targeted capture gene panels or less sensitive but more comprehensive genome sequencing. For example, several cases profiled using targeted gene capture assays yielded no therapeutically actionable alterations, possibly because the underlying causal mutations are not yet represented in the target lists. Likewise, at least one case of exome sequencing also yielded limited pathogenic information, presumably because of the relatively low tumor purity of the profiled specimen that was below the limit of detection of conventional exome capture coverage.

We also found that therapeutic recommendations required extensive, and in most cases, manual curation and analysis of the published literature. Although we used publically-accessible databases listing cancer gene mutations, such as canSAR [14], cBioPortal for Cancer Genomics [15, 16], and Tumor Portal [17], none of these repositories and annotation tools were found to have sufficient utility individually, requiring integration across several databases. Finally, in several cases, we identified alleles that are potentially pathogenic based on pathway and molecular modeling but that have not yet been reported to occur in tumor types under consideration, limiting the confidence of therapeutic recommendations based on such rationales.

In contrast to the previously described molecular tumor boards [7, 47, 48], we found a relatively high rate of adherence with the PTMB recommendations. Nearly half of all cases presented ultimately had alterations in clinical management based upon the PMTB recommendations. In prior reports, inability to access the desired targeted agents was cited as the most common reason for non-adherence to tumor board recommendations. [7, 8, 47, 48] We found only a single such instance for a patient in our study, who turned out to not meet eligibility criteria of a clinical trial.

It is possible that the largely relapsed and refractory nature of our patient cohort, and the absence of standard-of-care therapies for these groups of patients, contributed to the relatively frequent implementation of the PMTB recommendations. Only a minority of all pediatric patients treated at MSKCC were presented at the PMTB, presumably selected due to their clinically challenging or therapeutically refractory nature. As a result, patients reviewed by the PMTB had more advanced disease, more extensive prior treatment, and were older as compared to the overall MSKCC pediatric cohort. Thus, our study population had distinctive characteristics that may influence its generalizability.

Nonetheless, given our findings combined with other recently published experiences [7, 47, 48], we anticipate that molecular tumor boards will become increasingly used in pediatric oncology, at least in the near future. First, molecular and genomic tumor analysis yields complex, multi-variable profiles. It remains to be determined whether they will continue to require expert manual review such as the one implemented in our PMTB, or some aspects of the analysis can rely on rule-driven algorithms and formal clinical protocols, such as the NCI Molecular Analysis for Therapy Choice (MATCH) trial.

Second, we anticipate that the prospective evaluation of patients for tumor molecular profiling (as part of registries or by expert consultation) and selection of optimal molecular profiling technologies (targeted gene and transcript versus whole genome and transcriptome sequencing) will be needed to enable objective assessments of its clinical utility. It will be important to determine whether these assessments can be effectively accomplished using pre-treatment diagnostic biopsy specimens, or whether analysis of relapsed or recurrent tumors will be necessary, given the emerging evidence of drug response and tumor evolution that can affect clinical outcomes. [49, 50]

Increasingly, genome analyses are demonstrating that large subsets of patients have tumor mutations that are not highly prevalent in large unselected cohorts.[17, 51] Our study supports this notion, having identified novel mutations in specific tumor types with high likelihood of pathogenicity, though they have not been reported to be highly prevalent or even observed in certain cases. Thus, detailed data sharing and curation frameworks will need to be established to enable more accurate annotation of clinical molecular profiles. Such efforts are already being piloted, as in the AACR Project for Genomics, Evidence, Neoplasia, Information, Exchange (GENIE) that links clinical cancer genomic data across an international consortium. Finally, interventional clinical trials to determine the utility and effect of molecular profiling on clinical outcomes will be necessary.

## Acknowledgements

We thank all members of the Department of Pediatrics at MSKCC for their contributions to the PMTB, Alejandro Gutierrez, Marc Mansour, and Leo Wang for comments on the manuscript, and Daniel Baez for logistical support. This work was supported by the NIH K08 CA160660 (A.K.), the Burroughs Wellcome Fund (A.K.), and the Marie Josee and Henry R. Kravis Center for Molecular Oncology.

## Conflict of Interest Statement

The authors have no conflicts of interest to disclose.

